# Disturbance During Biofilm Community Succession Promotes Cooperation and Diversity

**DOI:** 10.1101/352914

**Authors:** James P. Stratford, Douglas M. Hodgson, Nelli J. Beecroft, Ann Smith, Julian R. Marchesi, Claudio Avignone-Rossa

## Abstract

Diversity-disturbance relationships have found widespread application in ecology, conservation and biodiversity management. In spite of their explanatory power, these conceptual frameworks have yet to be systematically applied to understanding succession in diverse microbial biofilms. Here we investigate community assembly in biofilms formed in replicate microbial bioelectrochemical systems using time-course sequencing of community 16S rRNA genes, corresponding to hundreds of operational taxonomic units (OTUs). For the first time we present a statistical model showing that a simple diversity-disturbance relationship can be used to explain dynamic changes in high diversity biofilm communities. This simple model reveals that succession in these systems is guided towards either a low diversity, generalist-dominated biofilm or a high diversity, cooperative-specialist biofilm, depending on the level of endogenous disturbance measured within the community. The pattern observed shows remarkable symmetry with findings from macro-scale communities such as grasslands, forests and coral reefs.

## Introduction

Mixed species biofilms can be comprised of thousands of bacterial taxa with a vast network of potential trophic interactions. Predicting the behaviour of these systems poses a similar challenge to that faced by ecologists studying macro-scale communities. Two of the most widely investigated ecological relationships are those between productivity and diversity as well as between disturbance and diversity. While some promising results have previously been obtained using simplified microcosms, thus far the operation of these relationships during succession within high diversity biofilm communities have not been detected (Konopka *et al*, 2015). The discovery of simple, familiar ecological patterns within diverse biofilms could provide a unique insight into the seemingly intractable complexity of mixed microbial community dynamics (Dini-Andreote *et al*, 2015).

A range of previous studies have shown clear relationships between community diversity and productivity (Grime, 1973a; Grime, 1973b; Gurevitch, 1986; Tilman *et al*, 1996; Grime, 1997; Buckling *et al*, 2000; Kassen *et al*, 2000; Gravel *et al*, 2011). The best known of these productivity diversity relationships (PDRs) is the unimodal or “hump backed” distribution (Grime, 1973). This predicts an increase in diversity with productivity which peaks at intermediate levels of productivity and then declines at high levels due to competitive exclusion. Disturbance diversity relationships (DDRs) are also mainstay conceptual frameworks in ecology (Connell, 1978; Wilkinson, 1999; Molino and Sabatier, 2001; Svensson *et al*, 2012). Disturbance has been shown to mitigate competitive exclusion, increasing diversity (Connell, 1978; Huston 1979) and cooperative interactions (Brockhurst *et al*, 2007). This is because competitive exclusion relies on the long-term exploitation of a small competitive advantage within a crowded system (Hautier et al, 2009). Disturbance damages established organisms, creating gaps which allow substitution with alternative community members. These gaps can also be exploited by less competitive organisms which may have greater rates of growth, dispersal or persistence (Tilman, 1994; Schnitzer and Carson, 2001). Disturbance can be exogenous or endogenous. Exogenous disturbance is caused by unique or external factors; in macro-scale systems this includes human activity or natural disasters. On the other hand, endogenous disturbance arises from many smaller routine events within the community (Peh *et al*, 2011). In macro-scale communities, sources of endogenous disturbance include forest tree-falls (Attiwill, 1993), herbivory and pathogens (Ayres and Lombardero, 2000), and may be induced by exogenous events such as forest clearing or climate change (Overpeck *et al*, 1990). Analogous processes in microbial communities are predation, phage infection, production of bioactive molecules (e.g. antibiotics) or biofilm disaggregation.

Advances in sequencing methods now permit characterisation of communities comprised of thousands of distinct bacterial taxa within a single analysis run (Caporaso *et al*, 2011). This is especially useful for microbial ecology, as large numbers of individual microbial taxa can be simultaneously detected based on their 16S rRNA gene. Therefore, in a time-course experiment, an entire community can be resolved into a comprehensive breakdown of operational taxonomic units and their relative abundances, as well as how these change with time. As a consequence, the fundamental rules governing community assembly should be detectable so long as other factors including inoculum, medium feed rate and biofilm substrate are kept constant. For comparison, the analysis for succession sequences in macro-scale communities requires labour spanning entire careers (Silvertown, 2006; Enquist *et al*, 2009).

Microbial fuel cells (MFCs) are bioelectrochemical systems (BES) that utilise the metabolic properties of bacterial biofilms to convert an electron donor, usually a carbon source, into carbon dioxide and hydrogen ions while transferring electrons to the anode on which they are growing, which acts as the final electron acceptor. Hydrogen ions migrate through a cation selective membrane towards the cathodic chamber, while electrons travel the external circuit to reduce an electron acceptor (e.g. molecular oxygen in air breathing systems) at the cathode (Figure 1). MFCs can be used as continuous biofilm culture systems by feeding the electron donor, usually a sugar, over extended periods of time (Logan *et al*, 2006; Beecroft *et al*, 2012; Stratford *et al*, 2014). In those systems, the electron transfer rate has been found to correlate strongly with CO_2_ production and with the decrease in chemical oxygen demand in the anode chamber (Thurston *et al*, 1984). Therefore, the electrical output of the system can be used as a convenient indicator of community productivity. The stoichiometry of the reactions required within the biofilm have been characterised in detail and assigned to distinct metabolic types (Freguia *et al*, 2008, Hodgson *et al*, 2016). In a carbohydrate-fed MFC (such as our system), the fuel is converted into fermentation products which are then consumed in electrogenic reactions mediated by anodophiles (Freguia *et al*, 2008; Kiely *et al*, 2011). This results in two broad types of community members, fermentative cells and anodophilic respirators, which correspond to the biochemical processes taking place within the biofilm. Organisms carrying out fermentation obtain metabolic energy by conversion of substrates into e.g. volatile fatty acids such as acetate, while anodophiles obtain their energy through the respiration of a carbon source, such as those fermentation products, using an anode as the final electron acceptor (Figure 1). Anodophiles must possess a respiratory electron transport chain in order to be electrogenic, and respiration is therefore a physiological necessity. Many bacterial species are generalists, capable of performing both respiratory and fermentative metabolism to obtain energy. These generalist species possess an electron transport chain in addition to the ability to ferment sugars. Bacteria can be identified and categorised as one or the other of these types: fermenter only (F), non-fermentative respirator (R), or respiratory fermenter (RF). Each of the alternative specialist types (R and F) requires the other to efficiently utilise the available resources, while the generalist species (RF) are potentially capable of powering the fuel cell without the need for cooperative syntrophy.

**Figure 1.**
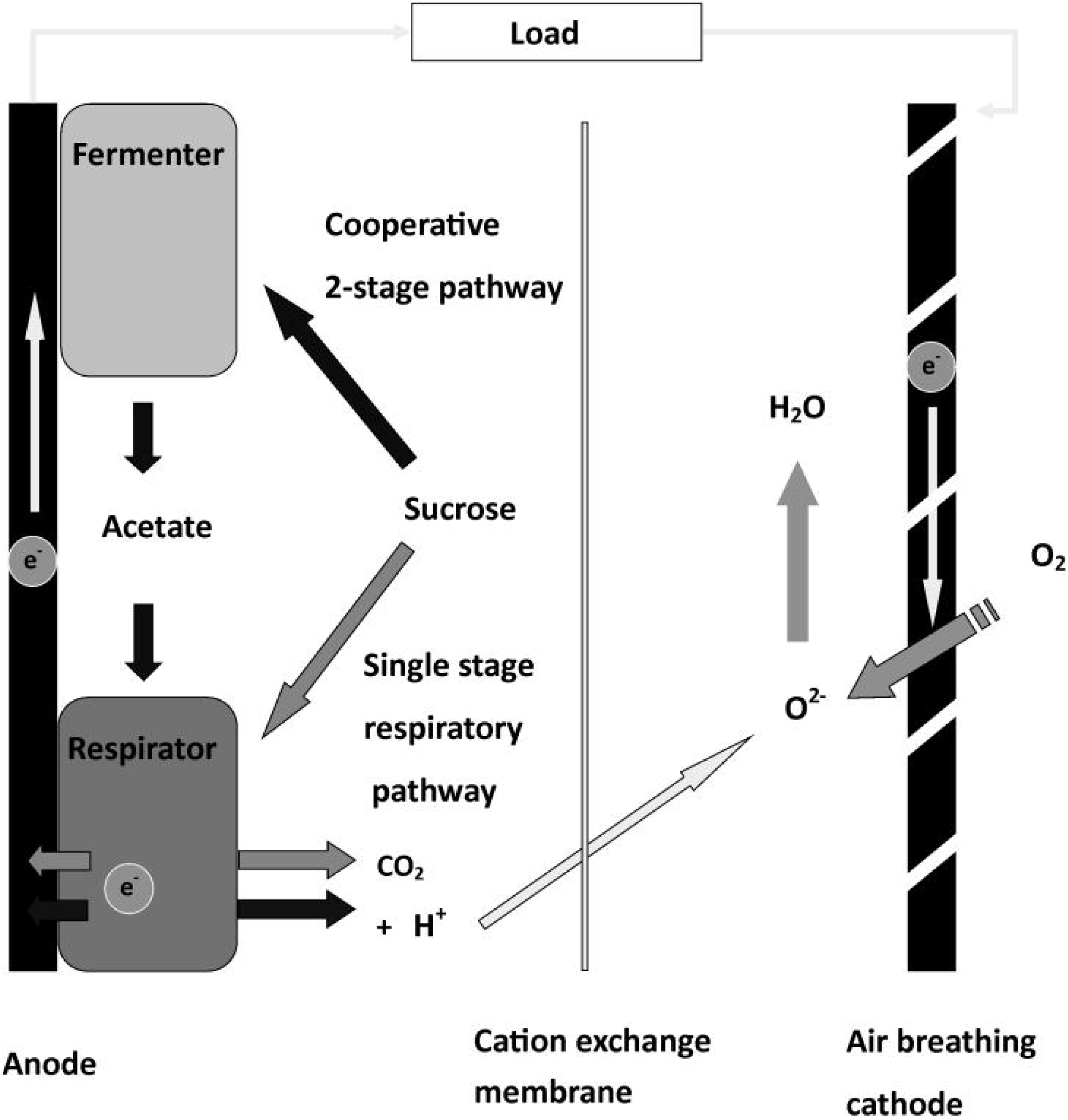
Simplified summary of current understanding of the biochemistry underlying general operation and syntrophy within MFCs.

We show here that microbial electrochemical systems can be used as a model for the balance between cooperative and competitive interactions within rich bacterial communities during community succession. Using microbial fuel cells as an experimental platform, we demonstrate that endogenous disturbance is associated with increased cooperation and diversity within multispecies bacterial biofilms. This relationship can also explain why diversity is greatest at intermediate levels of biofilm productivity.

## Results

Bacteria colonise the electrode rapidly and biofilms were established after 15 days. In spite of identical medium composition and reactor dilution rate, communities are highly dynamic and show substantial changes in composition with time. These changes occur not only through increases in sequence abundance but also sequence losses for many taxa. Some communities are more dynamic and experience larger losses whilst others are relatively stable. Community dynamics are most apparent from measurements of DNA content per area unit, with detectable DNA per cm^2^ increasing from a range of 48.1 - 99.7 ng μl^−1^ at 15 days to 97.1 – 154 ng μl^−1^ at 91 days.

Diversity values varied widely among reactors at a given time point and between different time points (Figure 2). At 15 days, diversity averaged 9.82 species equivalents while at 40 days this value was 8.08, and 9.66 at 90 days; overall, there was no significant trend across time, r= 0.09, p = 0.69. It is noteworthy that all communities had similar diversity at 15 days (8.12 - 12.14 species equivalents). Productivity increased with time, from a value of 456 mW m^−3^ at 15 days rising to 884 mW m^−3^ at 40 days and 1340 mW m^−3^ at 90 days. The relationship between productivity and diversity revealed a probability distribution in which all of the 5 most diverse communities were found to occur in the centre of the productivity range around 925 mW m^−3^ (Figure 3A).

**Figure 2.**
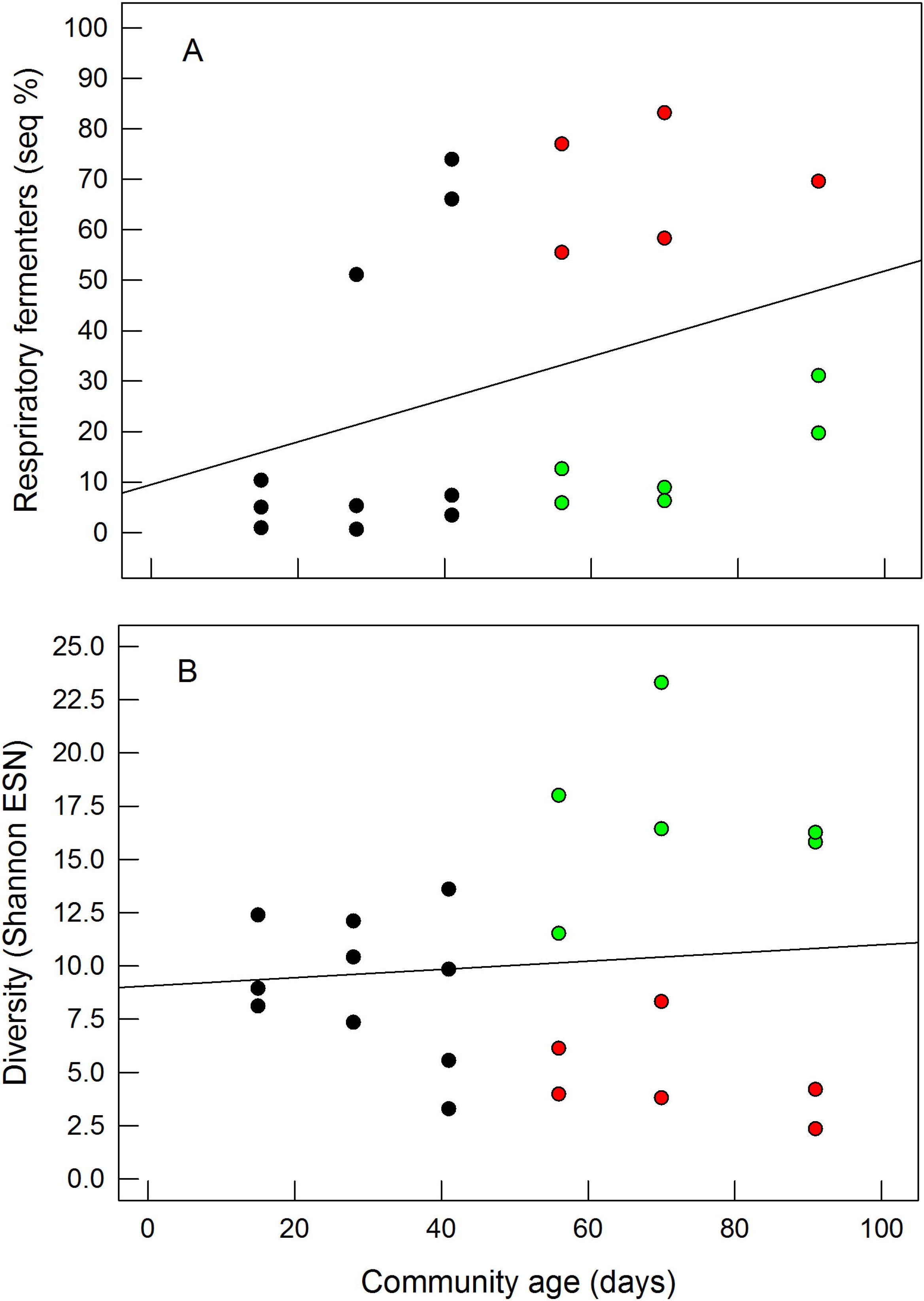
Divergent succession in electrogenic microbial communities; communities tend towards either generalist (RF) dominated or diverse (syntrophic mixture of specialist R and F bacteria). (A) Change in the proportion of generalist (RF) with time, r= 0.43, p = 0.049. (B) Change in Shannon diversity (units = equivalent species) with time, r= 0.09, p = 0.69. Open circles identify communities rich in generalists (RF) while closed grey circles identify communities with greater diversity. Symbol coding identifies the same communities in both (A) and (B).

**Figure 3.**
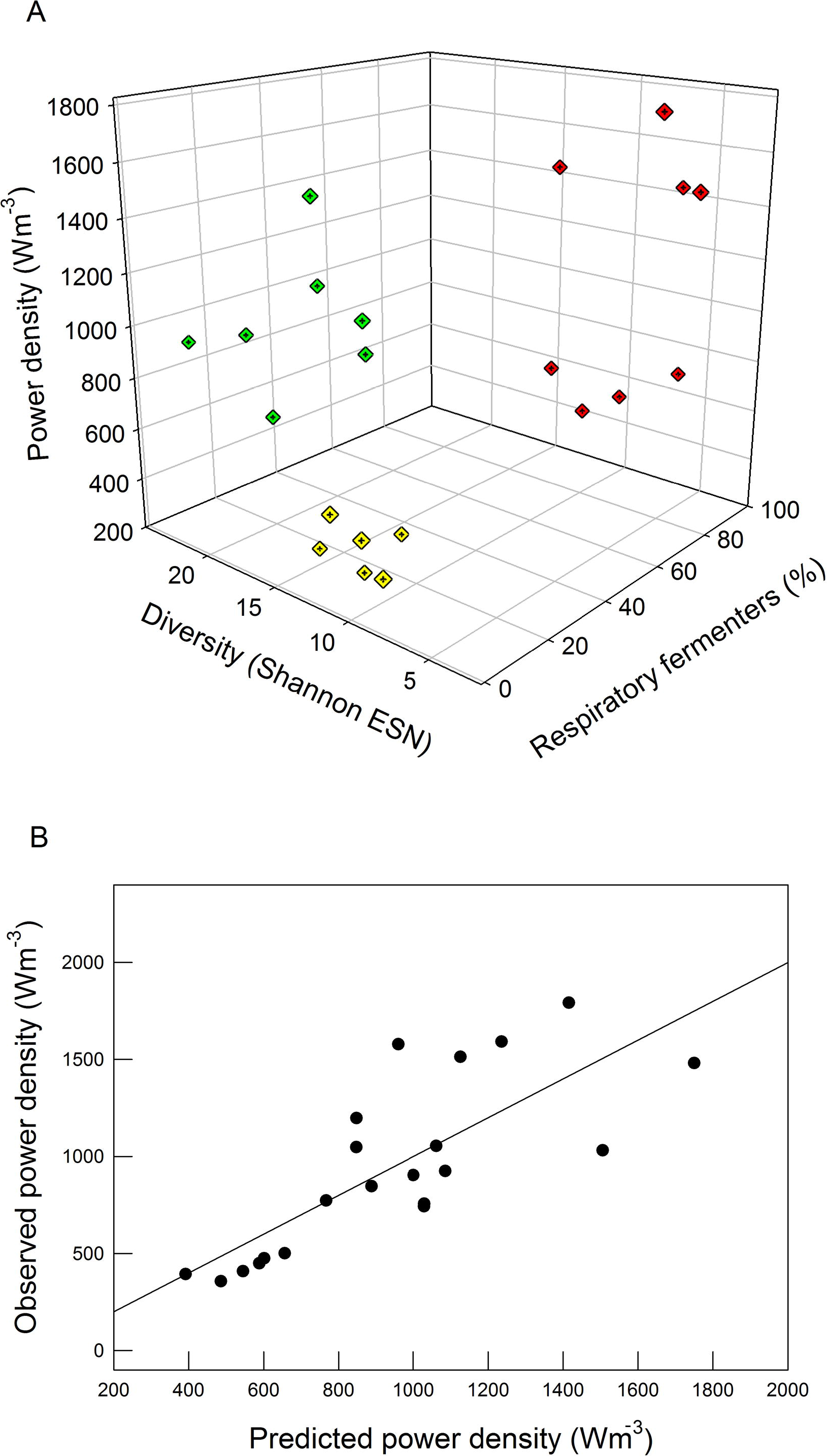
Alternative community profiles, either high diversity or generalist-dominant community types. (A) 3D graph simultaneously showing power density (absolute), Shannon diversity and relative abundance of RF (generalist) microbes. Generalists are defined as those organisms with the capacity for both respiration and fermentation (RF). (B) Plot showing a comparison between values of power density predicted by a simple linear model with the values observed for all communities. See Table 1 for linear model details.

**Table 1.** Summary for multiple linear model (MLM) predicting the productivity of MFCs. Power density per unit of detected community DNA was used as the dependent variable, while relative abundance of generalist taxa, DNA per cm^2^ of electrode and Shannon diversity were used as the independent variables.

**Multiple linear model summary**
|  |  |  |  |
| --- | --- | --- | --- |
| R | R<sup>2</sup> | R<sup>2</sup> adj | p |
| 0.75 | 0.56 | 0.48 | 0.003 |

|  | Coefficient | Std. Error | t | P |
| --- | --- | --- | --- | --- |
| Constant | - 9.422 | 7.814 | - 1.206 | 0.244 |
| Shannon Index | 9.289 | 2.757 | 3.370 | 0.004 |
| Generalist abundance (%) | 0.223 | 0.0526 | 4.236 | < 0.001 |
| DNA concentration (µg/cm<sup>2</sup>) | - 0.0688 | 0.0271 | - 2.539 | 0.021 |

The proportion of RF bacteria increased with time, r= 0.43, p = 0.049 (Figure 2A), and RF abundance positively correlated with MFC absolute power density, r = 0.55, p = 0.01 (regression not shown). This is clearly observed in the data presented in Figure 2: At later time points, higher abundance of RF is associated to a more productive community. RF abundance negatively correlated with community Shannon diversity, r = − 0.75, p < 0.001 (Figure 4). This mutually exclusive relationship appears as a bifurcation in the pattern of succession for both diversity and RF abundance, first appearing between 20 and 40 days (Figure 2). Combining data for generalist (RF) sequence abundance with Shannon diversity measurements (units = Shannon Index) allows the construction of a multiple linear model, R = 0.75, p = 0.003 (Table 1). The multiple linear model revealed positive relationships between the abundance of generalist (RF) bacteria and productivity as well as a positive relationship between diversity and productivity. This model explains 56% of all variance in power output between MFC communities. These relationships are integrated into a conceptual framework, summarised in Figure 6. Candidates for explaining this residual variation include fouling of the cathode or the ion exchange membrane (Zhuang *et al*, 2012), variation in the performance of different taxa, and stochastic variation. Most strikingly, the measured historical disturbance for a community predicts its succession towards either a high diversity community of specialists (R or F) or alternatively a low diversity generalist community. Disturbance (cumulative historical sequence loss) was strongly associated with community diversity, r = 0.81, p < 0.001 (Figure 5B). Partial regression between Shannon diversity and disturbance, controlling for community age, reveals an even stronger relationship, R = 0.89, p < 0.001 (regression not shown). This relationship demonstrates that cumulative disturbance with time (rather than community age) is more strongly associated with the observed trend. Disturbance also predicts generalist (RF) relative abundance, r = − 0.49, p = 0.037 (Figure 5A).

**Figure 4.**
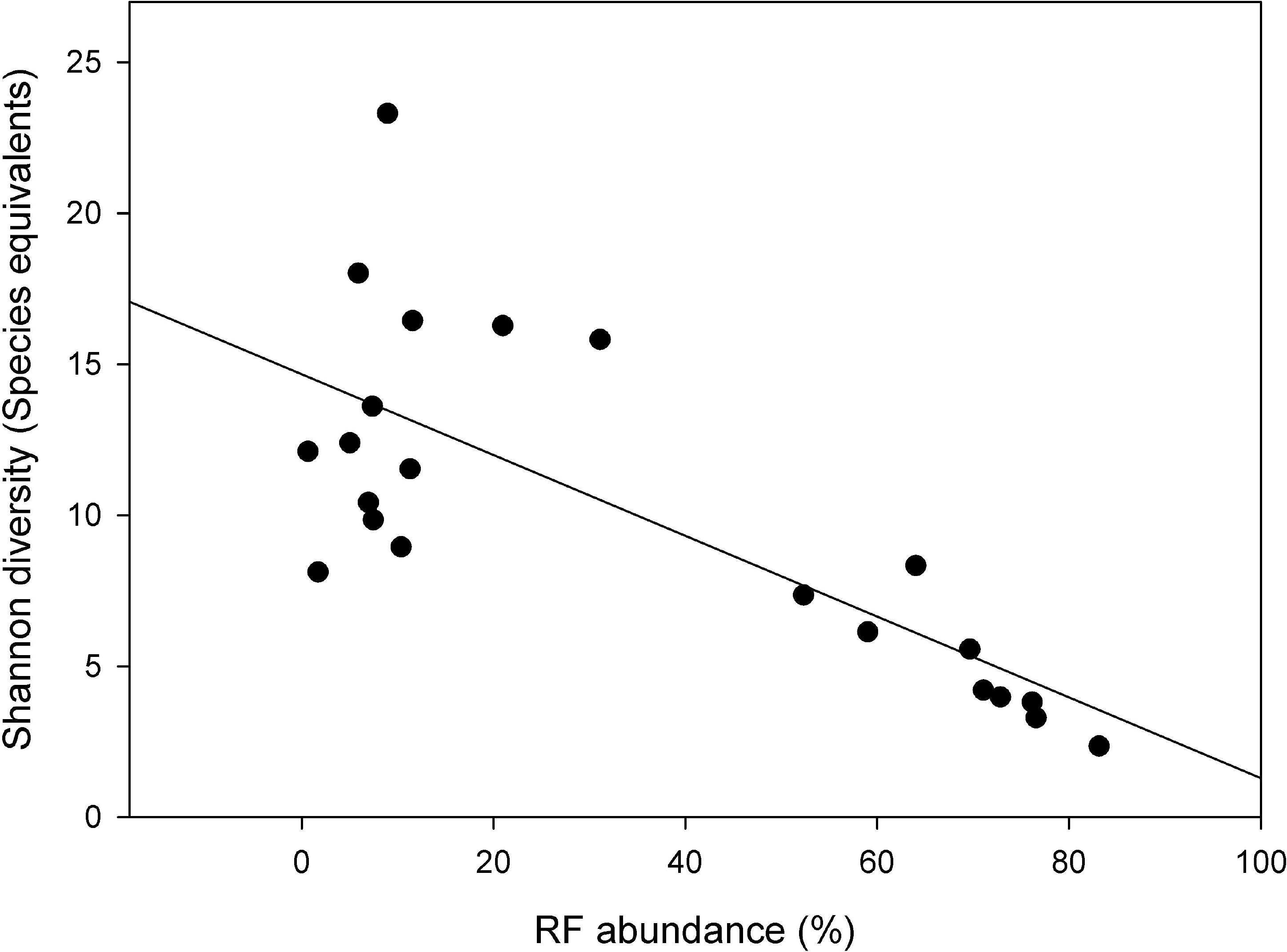
Relationship between the relative abundance of generalist and Shannon diversity in MFC communities, r = − 0.75, p < 0.001.

**Figure 5.**
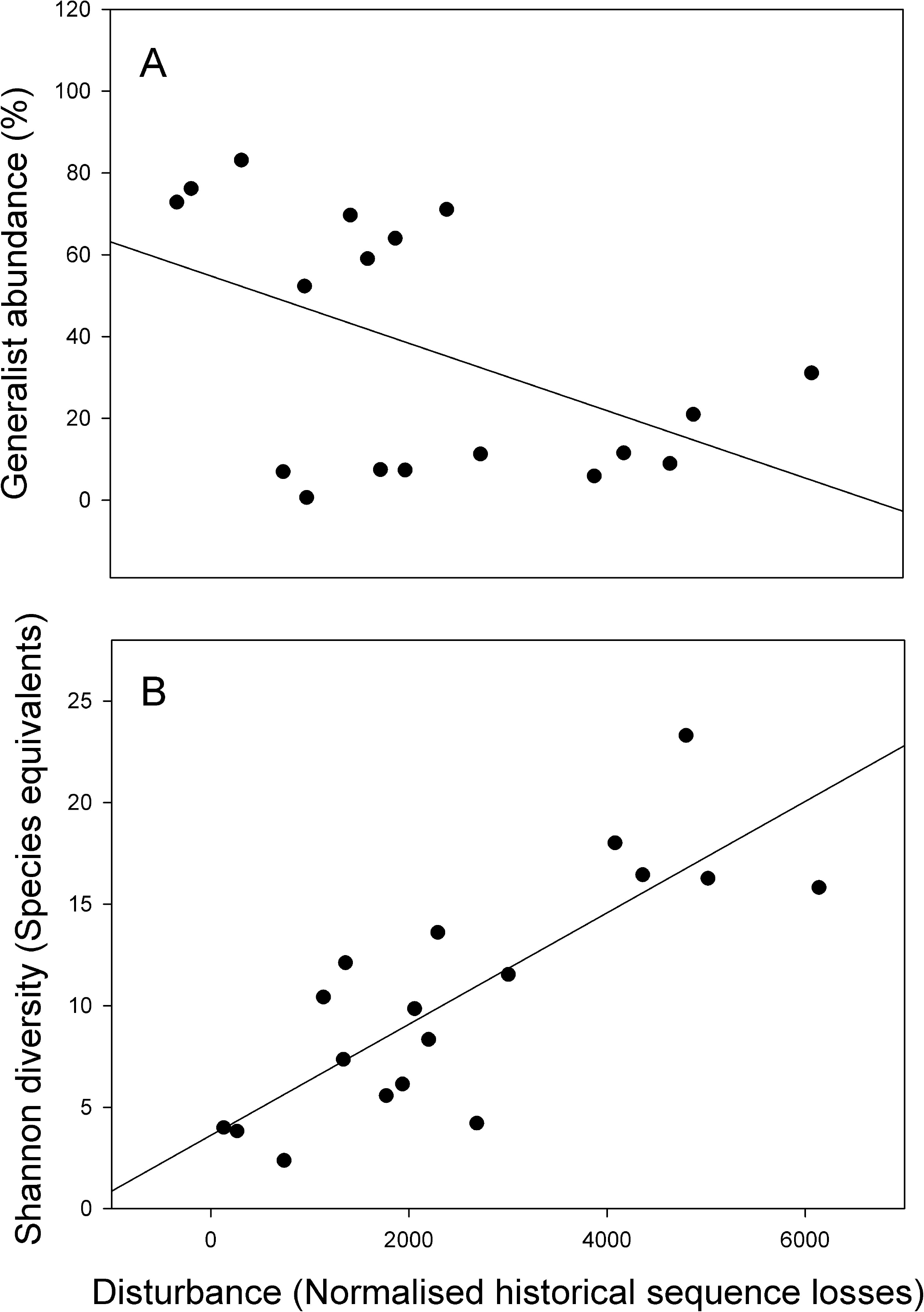
Relationship between historical disturbance in MFC communities and (A) generalist abundance, r = −0.49, p = 0.037. (B) Shannon diversity, r = 0.81, p < 0.001.

**Figure 6.**
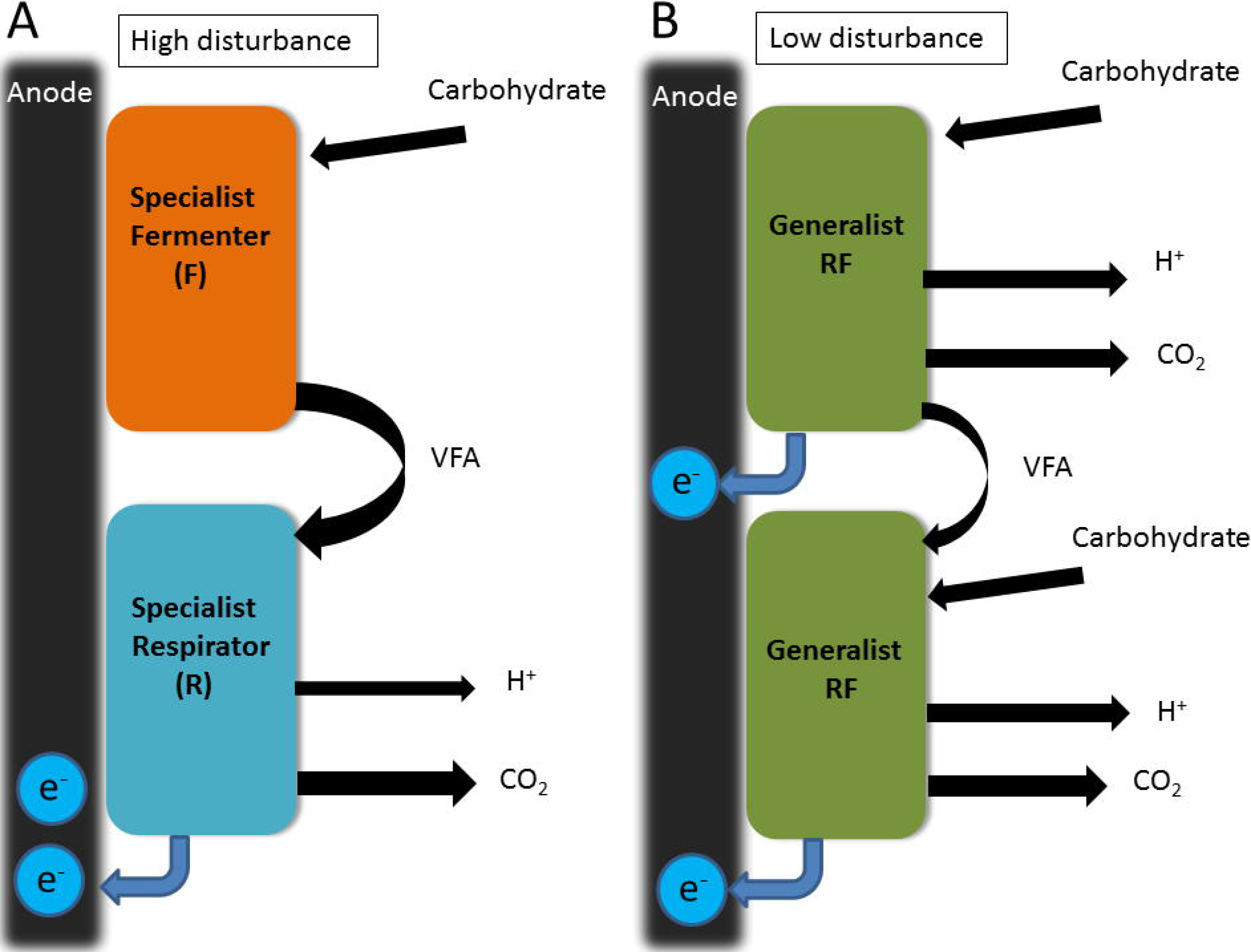
Schematic of alternative minimal community structures in MFCs under different disturbance regimes. Arrows indicate transfer of chemical species between biofilm cells.

## Discussion

In our model system, communities contained bacteria that exclusively utilised fermentative or respiratory metabolism, as well as generalist organisms capable of both metabolic processes (Figure 1) (Freguia *et al*, 2008, Hodgson *et al*, 2016). We have shown that both greater diversity within specialist communities and a high relative abundance of generalists increases electrogenic activity. As our purpose in this work was to detect simple patterns corresponding to those observed in macro-scale systems, we used the least complex statistical models which were sufficient to reveal these relationships. A multiple linear model (MLM) using both metrics as independent variables, while controlling for the amount of cells present (extractable DNA), explained a majority of the variation in MFC power density (Table 1). Where a generalist can function equally well within two niches, it is expected that it will have an advantage over specialists and be competitively dominant (Wilson and Yoshimura, 1994). This results in a diversity productivity relationship in which diversity is greatest at intermediate values of productivity. The operation of this principle in our system is strongly supported by the observation that all 5 of the highest diversity communities occur in the centre of the productivity range (Figure 3). This relationship is reminiscent of the hump-backed distribution discovered in other ecosystems (Grime, 1973; Huston, 1979). In line with the predictions of diversity disturbance relationships, disturbance was associated with reduction of the advantage possessed by generalists; and as a consequence, was associated with increased syntrophy between fermentative and respiratory bacteria. This diversity-disturbance relationship (DDR) coincided with community succession toward either a low diversity generalist or high diversity specialist community. We found in our system that the DDR has a monotonic pattern of increase in diversity with disturbance. This monotonic relationship is expected when the reproductive rates of all species exceed the maximum rate of mortality due to disturbance (Svensson *et al*, 2012). Under these conditions disturbance will never be high enough to reduce diversity, truncating the unimodal distribution. As most culturable bacteria have very short division times compared with our measurement intervals, the monotonic relationship would reasonably be expected for our system.

Synthetic systems have been suggested as experimental tools to control environmental constraints and reduce variability in microbial ecology studies (Konopka *et al*, 2015). Our results clearly demonstrate the potential for microbial electrochemical systems to be used as microcosms in which to model fundamental ecological relationships existing within microbial ecosystems, as it has been previously proposed (Dolfing 2014). Thus far, the bulk of microbial electrochemistry research is directed towards extending the range of feed-stocks, improving power output, optimising cell design and finding novel electrogenic bacteria (Jia et al, 2003; Logan *et al*, 2006; Chaudhury and Lovley, 2003; Yong *et al*, 2012; Zhuang *et al*, 2012). Microbial fuel cells generate a direct current which gives an instantaneous and easily measured proxy for community productivity (Thurston *et al*, 1984). The advent of next generation sequencing methods also allows the collection of detailed community abundance data for hundreds or thousands of species across replicate systems at multiple time points. This can be combined with the ability of MFCs to support high diversity biofilm cultures across extended time horizons, rendering anode associated electrogenic biofilms a tractable model for microbial community succession. This approach has allowed us to demonstrate the operation of a diversity disturbance relationship within electrode associated biofilms.

## Conclusions

A large amount of evidence has previously shown that diversity improves productivity in ecosystems (Tilman *et al*, 1996; Loreau *et al*, 2001). This corresponds to one possible trajectory within our Multiple Linear Model. Dominant organisms have also been shown to contribute more to system productivity than species with subordinate rankings (Smith and Knapp, 2003). This corresponds to the second possible trajectory within our 3D MLM. Our findings suggest that both of these relationships can function in MFC biofilm communities. Increased biodiversity improves community productivity for mixtures of specialists. However, a higher abundance of competitive dominants will also increase productivity. During succession, generalists (RF) will tend to prevail over specialist organisms and disturbance will tend to mitigate their advantage. Succession can follow either path from the same starting point and the trajectory taken depends on the level of disturbance measured. The alternative trajectories of succession we have observed, when examined together, explain the approximate unimodal distribution of diversity along the dimension of biofilm productivity. This pattern contributes to explaining variation in community structure where starting abiotic conditions are otherwise identical.

## Materials and Methods

### Microorganisms and media

MFCs were inoculated with anaerobic digester sludge originated in a biosolids mesophilic digester (Cog Moors Sewage Treatment Works, Cardiff, UK). Solids were removed by sieving through a 0.6-mm mesh (Endecotts Ltd., UK), and the collected material was stored at 4°C. The culture medium contained (g l^−1^): NH_4_Cl, 0.31; NaH_2_PO_4_·H_2_O, 5.38; Na_2_HPO_4_, 8.66; KCl, 0.13. The pH was adjusted to 7.0, and the medium was supplemented with 12.5 ml l^−1^ of a trace mineral solution and 12.5 ml l^−1^ of a vitamin solution (Lovley et al. 1984). The medium used for the batch operation contained 5 g l^−1^ sucrose, while the medium for the continuously operated MFCs contained 0.1 g l^−1^ sucrose. All media were autoclaved at 121°C for 15 min, except for the vitamins, mineral and sucrose solutions, which were sterilised by filtration through a 0.2-μm pore size membrane (Nalgene, USA).

### MFC set up

MFCs consisted of 9 cm^3^ Perspex anode chambers and cover plates, with stainless steel metal plates serving as a contact between the air-breathing cathode and the electrical circuit. The anode (geometric area: 32 cm^2^) was made of carbon fibre veil (PRF Composite Materials, UK) with polyvinyl alcohol binder to improve anodic capacity. The anode was connected to the electrical circuit with an insulated Ni/Cr wire weaved across the anode. The air-breathing cathode was made of type A carbon cloth (geometric area 9 cm^2^; E-TEK, USA) coated with 4 mg cm^−2^ of Pt black catalyst and polytetrafluoroethylene binder. The Pt side of the cathode was painted with 0.5– 1.0 mg cm^−2^ of Nafion perfluorinated ion-exchange ionomer (5% w/v in lower aliphatic alcohols and H_2_O). Anode and cathode chambers were separated by a Nafion-115 proton-exchange membrane (20 cm^2^, DuPont, USA). The membrane was pre-treated by boiling sequentially for 1 h in 6% w/v H_2_O_2_, H_2_O, 0.5 M H_2_SO_4_ and H_2_O. The pre-treated PEM was stored in deionised water sheltered from light until assembling the MFC.

### MFC operation

Replicate MFCs were inoculated with a 10% v/v suspension of anaerobic digester sludge in sucrose-containing medium. MFCs were operated for approximately 2 weeks to ensure establishment of the anodic biofilm. During that time, the anodic suspension was replaced with fresh N_2_-purged fresh medium until a stable biofilm was present, as assessed by constant output and repeatable cycles of voltage generation. The MFCs were then operated in continuous mode, supplying fresh medium at a flow rate of 0.18 mL min^−1^. The MFCs were operated at room temperature (21–22°C), and anaerobic conditions were kept by maintaining a continuous flow of oxygen-free N_2_. Samples were taken for chemical and microbial community analysis.

### Chemical analyses

Total carbohydrate consumption was calculated as the difference between the concentration of carbohydrates in the influent and the effluent medium, determined by a colorimetric method (Dubois et al. 1956). Chemical oxygen demand (COD_Cr_) was analysed according to the standard method (SFS 5504 1988). Coulombic efficiency was calculated as previously described (Logan et al. 2006). The pH of the effluent medium was monitored using a pH meter (Mettler Toledo MP220, Switzerland).

### Electrochemical measurements

The electrical output was monitored using a battery tester (Arbin BT2000, Arbin Instruments, USA) controlled by dedicated software (MITS Pro, Arbin Instruments, USA) across a fixed external resistance of 40 kΩ. Polarisation curves were recorded by measuring the decrease in voltage when external resistances were varied over the range 700 kΩ to 500 Ω, for 5 min for each resistance. The volumetric power density was calculated as P=UI/V, where U is the measured voltage, I is the current and V is the volume of the anodic suspension. The ohmic internal resistance of the MFCs was measured by electrochemical impedance spectroscopy. Impedance spectra were recorded between anode and cathode in the frequency range of 0.1 Hz– 1 MHz and with a sinusoidal perturbation of 10 mV amplitude under open circuit voltage using a frequency response analyser (Solartron Analytical 1260) and a potentiostat/galvanostat (Solartron Analytical 1287, Solartron Analytical, UK) (Zhao et al. 2008).

### Microbial community analysis

Total DNA samples were obtained from the anodic biofilm and from the anodic suspension using FastDNA Spin Kit for Soil (MP Biomedicals, UK). MFCs were temporarily disassembled in an aseptic environment, and a 1 cm^2^ fragment of the anode was cut out with a sterile scalpel. The MFCs were immediately reassembled and operation was restarted. For the planktonic DNA samples, anode suspension samples were centrifuged (10,000×g, 5 min), washed three times with 1 ml phosphate-buffered saline (NaCl, 8.0 g.l-1; KCl, 0.2 g.l-1; Na2HPO4, 1.15 g.l-1; KH2PO4, 0.2 g.l-1; pH 7.3) and resuspended in 100 μl of nuclease-free water (Promega, UK).

### Sequence analysis

Fifty-four samples were analysed in Mothur using the 454 SOP Pipeline (Langille et al 2013). The forward primer used was 28F-GAGTTTGATCNTGGCTCAG. To reduce sequence error, trim.seqs() was implemented, with the parameter minlength set to 250. The kmer searching method was employed to align sequences by using a Silva bacterial database (www.arb-silva.de/), with the flip parameter set to true, allowing for the reverse complement of the sequence to be aligned for better results. The screen.seqs command was implemented to remove any sequences outside the 2.5%-tile to 97.5%-tile range of sequences. The filter.seqs was used to remove empty columns from the alignment, which gives the length of filtered alignment to 897. To classify the sequences, we used a RDP database/reference sequence files and the Wang method (Wang et al 2007).

To minimise artefacts and maintain the same number of reads in each sample, singleton OTUs and OTUs < 10 reads in any sample were collated into OTU_singletons and OTU_rare phylotypes, respectively. The taxonomic affiliation of the partial 16S rRNA gene sequences (from Phylum to Genus) were determined using the RDP MultiClassifier script (Wang et al 2007) to generate the RDP taxonomy while species level taxonomies of the OTUs were determined using the USEARCH algorithm (Edgar 2010) combined with the cultured representatives from the RDP database. Alpha and beta indices were calculated from these datasets with Mothur and R using the Vegan package (Oksanen et al 2013).

PICRUSt analysis allows for phylogenetic prediction of organismal traits using the 54 samples from the Mothur analysis. The shared file (representing the number of times that an OTU is observed in multiple samples) generated by Mothur was converted into a biom file (make.biom), from where it is possible to follow the PICRUSt metagenomics pipeline (Langille et al 2013). The first step was the normalisation of the OTU table (biom), enabling the creation of a metagenomics functional prediction table (predict_metagenomes.py). Using -- type_of_prediction parameter, allows for KEGG Orthologs predictions and COGS analysis on the samples. We used the collapse predictions into pathways function (categorize_by_function.py) to examine KEGG results from a higher level within the pathway hierarchy.

### Statistical analysis and model construction

Sequencing data was used to calculate the Shannon index, a measure of the diversity of the biofilm microbial communities. This can also be expressed as a true diversity value with units of equivalent species (ES) (Hill, 1973; Stratford *et al*, 2014). We classified the OTUs identified in the anodic biofilms into metabolic types based on their known metabolism or that of a closely related species. The metabolic types R (non-fermentative respirator), F (fermenter) and RF (respiratory fermenter) were assigned to taxa comprising 99% of total community sequences. Abundance values used in statistical analysis represent relative abundance of sequences attributed to organisms presenting the particular metabolic type in a survey of the literature. Raw data and details of the references used for classification are provided in the supplementary material. The effect of endogenous disturbance in bacterial biofilms was estimated from changes in the biofilm population. While it is currently impossible to detect individual disturbance events or mortality on an individual cellular level in microbial communities, an estimate can be calculated by measuring total sequences lost between measurement intervals and normalising these losses to the change in total sequences detected. Sequence losses measured in this way provide an estimate for disturbance in the time interval between measurements. The models built here are robust to the use of either the raw loss in sequences or the normalised value. Disturbance may also be interpreted as an indication of other dynamic processes within the system, e.g. cell removal from the biofilm by localised disaggregation or similar mechanisms not constituting mortality. These definitions of disturbance have previously been used interchangeably (Brockhurst *et al*, 2007) and are used in this context for our analysis. Here we define cumulative disturbance as the sum of all losses in the abundance of individual taxa between sampling points measured up to a given time point. In our system, all MFCs were inoculated with the same starting community and treated consistently thereafter, minimising externally induced variation in biofilm disturbance between communities. This means that losses within the microbial community should be predominantly due to endogenous disturbance rather than operator intervention. No assumptions have been made regarding the source and nature of disturbance during construction of our quantitative models. All correlations and regressions were carried out using SigmaPlot v.12.3 (Systat Software, San Jose, CA).

## Acknowledgements

CAR, DH, AS, JRM and MEB were supported by grants BB/J01916X/1 and BB/J019143/1 from the Biotechnology and Biological Sciences Research Council (BBSRC), under the Integrated Biorefining Research and Technology Club (IBTI) initiative. NJB and CAR were partially supported by the Engineering and Physical Sciences Research Council (EPSRC) Supergen 5 Biological Fuel Cells Consortium (Grant EP/H019480/1). JPS was funded by the BBSRC/EPSRC Synthetic Biology Centre grant BB/M017982/1 to the University of Warwick.

The authors declare no conflict of interest.

